# Liver-Resident Metabolic Reprogramming and Th2/1 Cell Accumulation Drive Anti-Helminth Immunity During *H. bakeri* Infection

**DOI:** 10.64898/2026.09.21.752666

**Authors:** Joshua Adjah, Zaneta D. Musimbi, Ankur Midha, Susanne Hartmann, Sebastian Rausch

## Abstract

The liver plays a pivotal yet understudied role in anti-helminth immunity. Here, we reveal that during *Heligmosomoides bakeri* infection, the liver serves as a critical immunological site, accumulating Th2 and Th2/1 hybrid cells through CXCR3-mediated recruitment driven by IFN-γ and the chemokines (CXCL9/10/11). Unlike traditional lymphoid organs, the liver maintains these functional Th2 and Th2/1 cells throughout the chronic and memory phases of infection, exhibiting potent cytokine production and no signs of exhaustion. Transcriptomic analysis of liver tissues uncovered stage-specific metabolic reprogramming, with early *H. bakeri* infection (6 days p.i.) suppressing oxidative phosphorylation and acute infection (14 days p.i.) activating immune pathways (e.g., TNF, JAK-STAT). Notably, metabolic shifts were independent of T cell infiltration and were instead induced by parasite excretory-secretory products (HES) and alarmins, which disrupted mitochondrial ATP production. The liver’s glucose-rich microenvironment supported T cell effector function, including IL-5 production linked to elevated serum IL-5 and bile IgA. Our findings redefine the liver as a dynamic regulator of anti-helminth immunity, integrating metabolic and immune responses to sustain long-term host defense. These insights open new avenues for targeting liver-specific pathways to enhance parasite clearance.

## Introduction

*Heligmosomoides bakeri* (*Heligmosomoides polygyrus*) is a naturally occurring intestinal nematode of rodents, parasitizing the small intestines of house mice and deer mice across Europe and North America (Behnke & Harris, 2010). *H. bakeri* establishes chronic infections and serves as a key laboratory model for studying immune responses, toxicology, and zoonotic nematode epidemiology (Behnke et al., 2009). Its ability to persist in immunocompetent hosts underscores its immunoregulatory prowess, making it a valuable analog for human hookworm infections and veterinary helminthiases (Behnke & Harris, 2010).

The gut-liver axis has gained prominence for its role in bidirectional immune crosstalk, with the liver acting as the first line of exposure for gut-derived antigens (Zheng & Wang, 2021; Albillos et al., 2020). While the liver is traditionally viewed as tolerogenic due to constant exposure to dietary and microbial antigens and as a site of exhausted T cell depletion (Crispe et al., 2000), our and other data challenge this paradigm (Ghilas et al., 2020; Pallett et al., 2017; Pallett & Maini, 2022). We recently reported significant enlargement of liver-draining lymph nodes (LLNs) in *H. bakeri*-infected mice, suggestive of an active hepatic involvement in anti-helminth immunity, potentially facilitating parasite clearance (Adjah et al., 2025). This aligns with documented immune cell dissemination to the liver during helminth infection (Mohrs et al., 2005), highlighting the need to explore how the liver and hepatic lymphoid tissues shape systemic responses.

Central to this antihelminth immune response are Th2/1 hybrid cells, a plastic T helper subset co-expressing Th2 (GATA3) and Th1 (T-bet) markers (Affinass et al., 2018; Burt et al., 2022). These hybrid cells arise during helminth infections in mice and humans (Bock et al., 2017), producing both type 2 cytokines and IFN-γ, although in lower proportions than Th1 and Th2 cells, and navigate complex immune challenges (Peine et al., 2013). In *H. bakeri* infection, Th2/1 cells populate the spleen, peripheral blood, and small intestines, expressing gut-homing receptors (CCR9, α4β7) akin to Th2 cells (Adjah et al., 2024). Strikingly, their abundance inversely correlates with impaired parasite clearance: C57BL/6 mice, which develop age-dependent Th2/1 expansions, fail to expel *H. bakeri* as efficiently as strains with fewer hybrids (Kapse et al., 2022). This suggests that Th2/1 cells may dampen protective Th2 responses and act as systemic confounders.

Regulatory T cells (Tregs) add a further layer of control, expanding in gut-draining tissue during *H. bakeri* infection to restrain rather than diversify Th2 immunity, partly via parasite-derived products that mimic TGF-β (Grainger et al., 2010; Finney et al., 2007). Their relationship with protection is non-linear, however: partial Treg depletion boosts Th2 output and clearance, yet complete removal paradoxically worsens worm burden through unchecked, damaging IFN-γ-driven immunopathology (Rausch et al., 2009; Smith et al., 2016).

These IFN-γ-producing cells localize to extra-mucosal sites, potentially sustaining systemic IFN-γ levels in a type 2-skewed environment (Affinass et al., 2018; Kapse et al., 2022). Meanwhile, LLN enlargement during infection suggests that hepatic lymphoid tissue actively shapes immune outcomes, though its role in promoting Th2/1 hybrids or Tregs remains unclear. We, however, observe that the liver harbors the highest proportions of highly functional (in terms of cytokine production) Th2/1 cells alongside Th2 cells compared with all previously studied organs.

We hypothesize that the liver dynamically modulates T cell responses during *H. bakeri* infection, influencing parasite clearance and systemic immunity. To test this, we tracked local and systemic T helper cell responses across infection stages, larval gut development (6 d.p.i), life cycle completion (14 days), and chronic phases (> 56 days), to dissect how host factors and organ-specific interactions dictate infection outcomes. Our study reveals that the liver is a key site of anti-helminth immunity, recruiting and maintaining Th2/1 hybrid cells via CXCR3/IFN-γ signaling during *H. bakeri* infection. Unlike lymphoid organs, the liver sustains these functional T cells long-term, supported by stage-specific metabolic reprogramming triggered by parasite-derived factors and the liver’s glucose-rich environment. By integrating the gut-liver axis and Th2/1 hybrid cell biology, this study aims to redefine the liver’s role in helminth immunity and uncover novel therapeutic targets for chronic nematode infections.

## Results

### Systemic T Cell Redistribution and Liver Enrichment of Th2/1 Cells During *H. bakeri* Infection

Consistent with reports of systemic T cell dissemination during *H. bakeri* infection (Mohrs et al., 2005; Classon et al., 2022), we observed T cell redistribution from lymphoid to non-lymphoid compartments, prominently to the liver (Figure 1A). By day 6 post-infection, GATA-3+ T cells were distributed across MLNs, LLNs, spleen, blood, and siLP, but interestingly the liver harbored a higher proportion than the siLP despite its distance from the infection site. By day 14, GATA-3+ T cells increased >3-fold in the gut and further accumulated in the liver, where Th2/1 cells (T-bet+GATA-3+) comprised nearly one-third of hepatic T cells at both time points (Figure 1A, B, C). While lymphoid compartments had minimal Th2/1 cells, the spleen, blood, and siLP showed detectable levels, with liver Th2/1 populations expanding by day 14 (Figure 1C). Effector T cells in the spleen and liver exhibited high co-expression of IL-4 and IFN-γ compared to other tissues (Figure 1D). Upon PMA/ionomycin stimulation, liver cells produced robust Th2 cytokines (IL-5, IL-13; Figure 1D, E), but low IL-10 within IL-4+ populations, unlike lymphoid compartments (MLN, LLN), which showed elevated IL-10, suggesting regulatory activity (Figure 1E). These findings identify the liver as a major site of GATA-3+ T cell accumulation, enriched in Th2/1 hybrids with dual Th2 (IL-4/5/13) and IFN-γ activity during infection.

**Figure 1:**
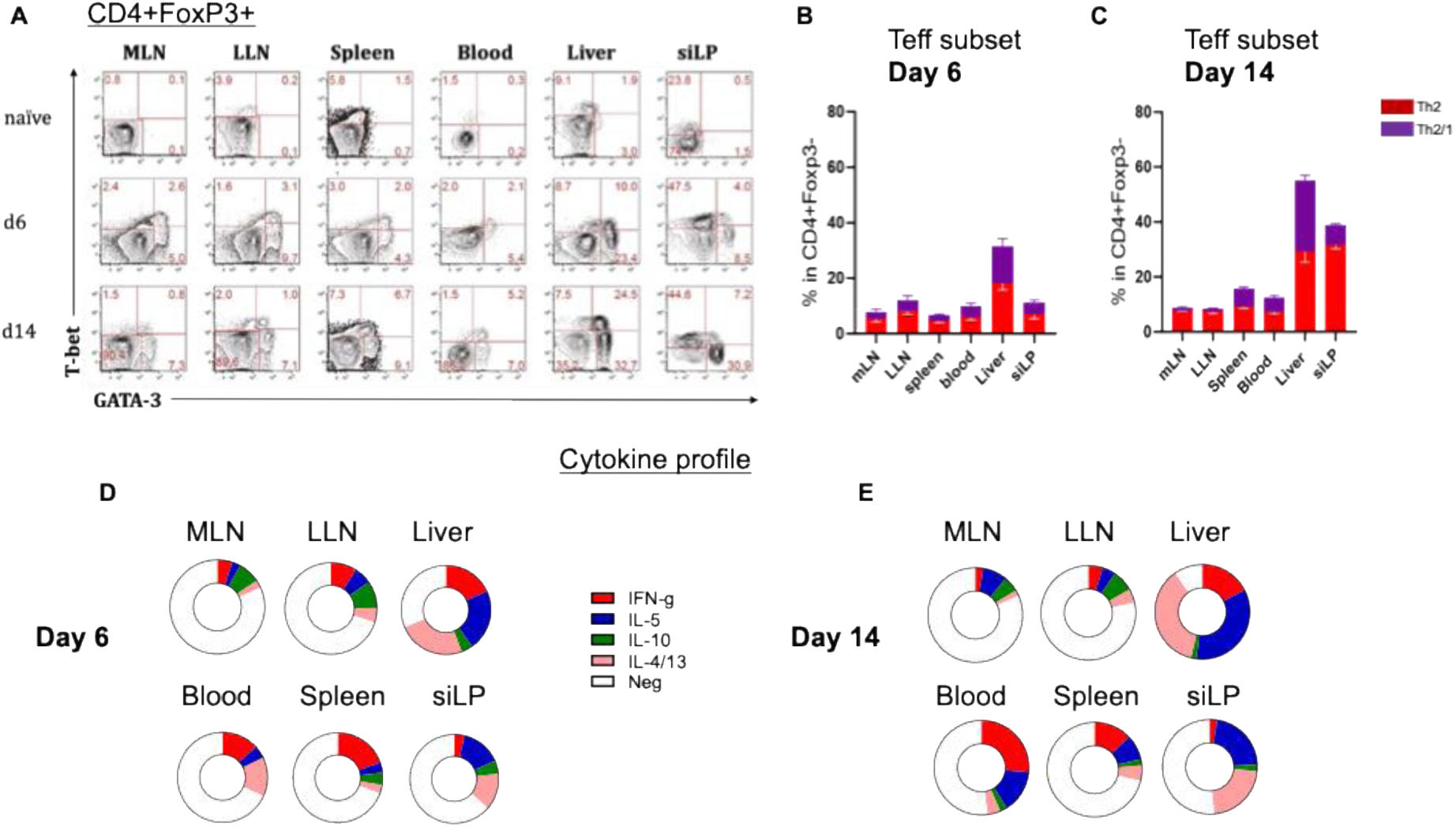
Organ-specific distribution and cytokine profiles of Th2 and Th2/1 hybrid cells during *H. bakeri* infection. (A) Flow cytometry plots showing T-bet and GATA-3 expression in CD4⁺FoxP3⁻ T cells from the indicated tissues in naïve mice and at days 6 and 14 post-infection. (B, C) Frequencies of Th2 (red) and Th2/1 (purple) subsets among CD4⁺FoxP3⁻ cells at day 6 (B) and day 14 (C). Bars represent mean ± SEM. (D, E) Pie charts summarizing cytokine expression profiles of CD4⁺ effector T cells across tissues at day 6 (D) and day 14 (E). Data are from ≥2 independent experiments. MLN, mesenteric lymph nodes; LLN, liver-draining lymph nodes; siLP, small intestinal lamina propia.

### Liver as a Reservoir of Functional Th2/1 Cells During Chronic, Memory and Challenge *H. bakeri* Infection

Contrary to suggestions that anergic T cells home to the liver for clearance (Zheng & Tian, 2019), we found hepatic T cell accumulation during *H. bakeri* infection represents sustained, functional populations. Analyzing C57BL/6 mice at chronic (day 56), memory (2 weeks post-cure), and challenge (3 days post-reinfection) phases revealed that while siLP and PEC contained conventional Th2 cells, the liver maintained comparable GATA-3+ T cell proportions but was dominated by Th2/1 hybrids, a pattern sustained through memory and expanded after challenge (Figure 2A-C). The spleen showed minimal Th2 involvement.

**Figure 2:**
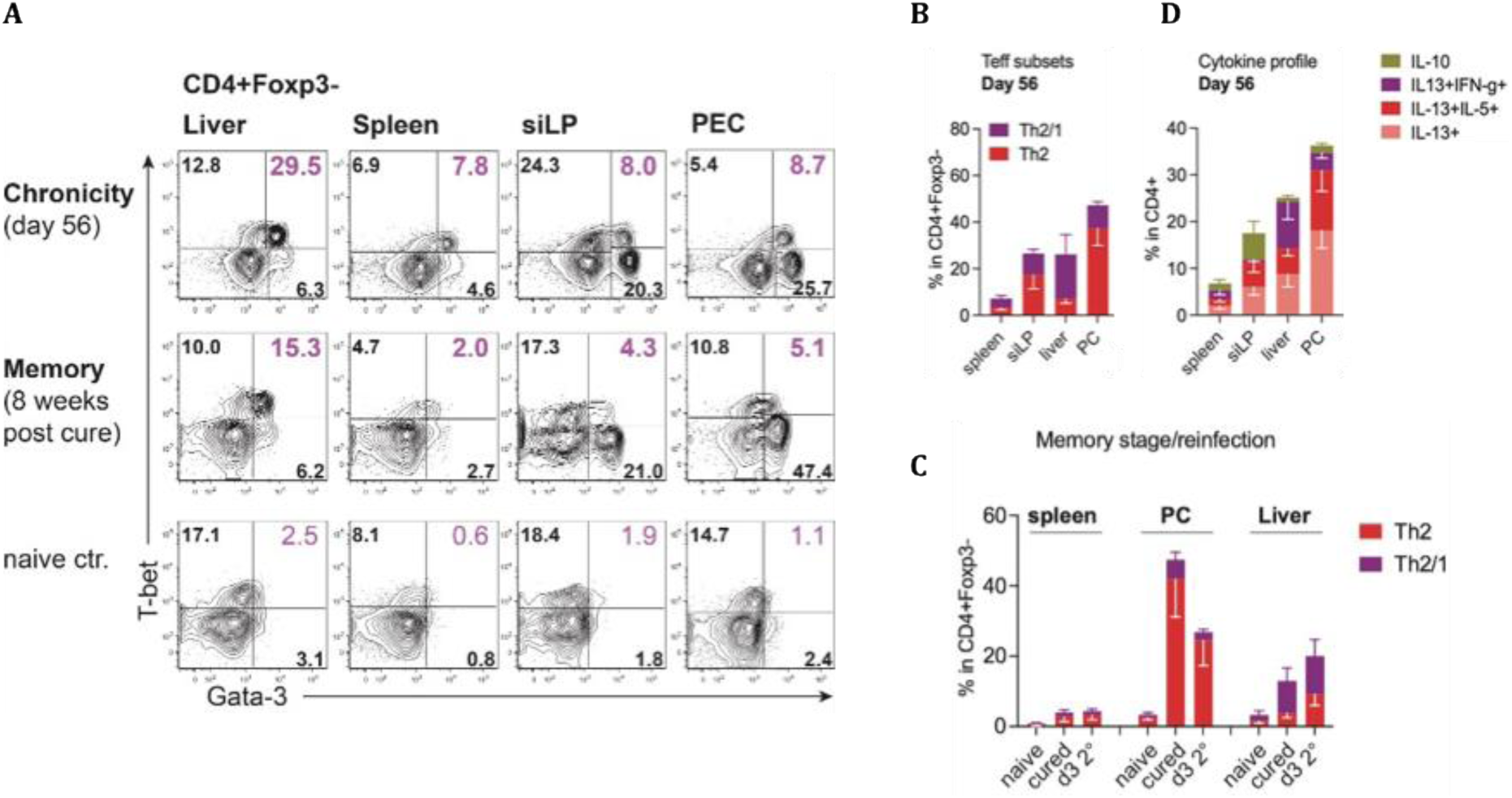
Tissue-specific dynamics of CD4⁺Foxp3⁻ T cell subsets during chronic and memory phases of *H. bakeri* infection. (A) Representative flow plots of T-bet+ and GATA-3+ T cell levels in liver, spleen, siLP, and PEC during chronic infection (day 56) and the memory phase (8 weeks post-cure). (B) Quantification of Th2 (GATA-3⁺T-bet⁻) and Th2/1 (GATA-3⁺T-bet⁺) subsets across tissues at day 56 post-infection. (C) Th2 and Th2/1 subset frequencies in spleen, PEC, and liver during memory phase and after reinfection. (D) Cytokine profiles of CD4⁺ T cells at day 56, showing percentages of IL-10⁺, IL-13⁺IFN-γ⁺, IL-13⁺IL-5⁺, and IL-13⁺ cells. Data represent mean ± SD (B, C, D).

Functional profiling showed liver-specific enrichment of IL-13+IFN-γ+ Th2/1 cells upon stimulation, while PEC cells produced predominantly Th2 cytokines (Figure 2D). The liver exhibited the lowest IL-10 expression, whereas siLP cells showed high IL-10, suggesting exhaustion (Figure 1D and E, 2D). Proliferation (Ki67) and apoptosis (Annexin V) assays indicated liver T cells were highly active, while siLP cells were anergic in the memory phase but recovered post-challenge, implying extra-intestinal replenishment (Supplementary Figure 1).

To determine if the liver replenishes exhausted siLP T cells, we assessed gut-homing marker expression (CCR9/α4β7) (Adjah et al., 2025). Liver T cells lacked these markers, implicating PEC rather than liver as the source of siLP replenishment (Supp. Figure 1). These findings position the liver as a persistent reservoir of functional Th2/1 cells, distinct from Th2-dominated siLP and PEC, with PEC potentially supporting intestinal repopulation during reinfection.

Our data demonstrate that highly functional GATA-3+ T cells persist in the liver throughout infection phases, with notable Th2/1 enrichment compared to spleen, siLP, and PEC. While siLP and PEC T cells show low proliferation, PEC cells are not anergic or exhausted, whereas siLP cells display anergy or exhaustion during the memory phase (Supplementary Figure 1).

### IFN-γ and CXCR3 Signaling Drive Liver-Specific Th2/1 Cell Accumulation

To identify drivers of Th2/1 accumulation, we assessed organ-specific IFN-γ, given the cells’ IFN-γ competence (Peine et al., 2013). The liver had significantly higher baseline IFN-γ+ CD4+ T cells than MLNs, LLNs, and spleen (Figure 3A), correlating with Th2/1 enrichment (Figure 3F), suggesting IFN-γ promotes hepatic localization.

**Figure 3:**
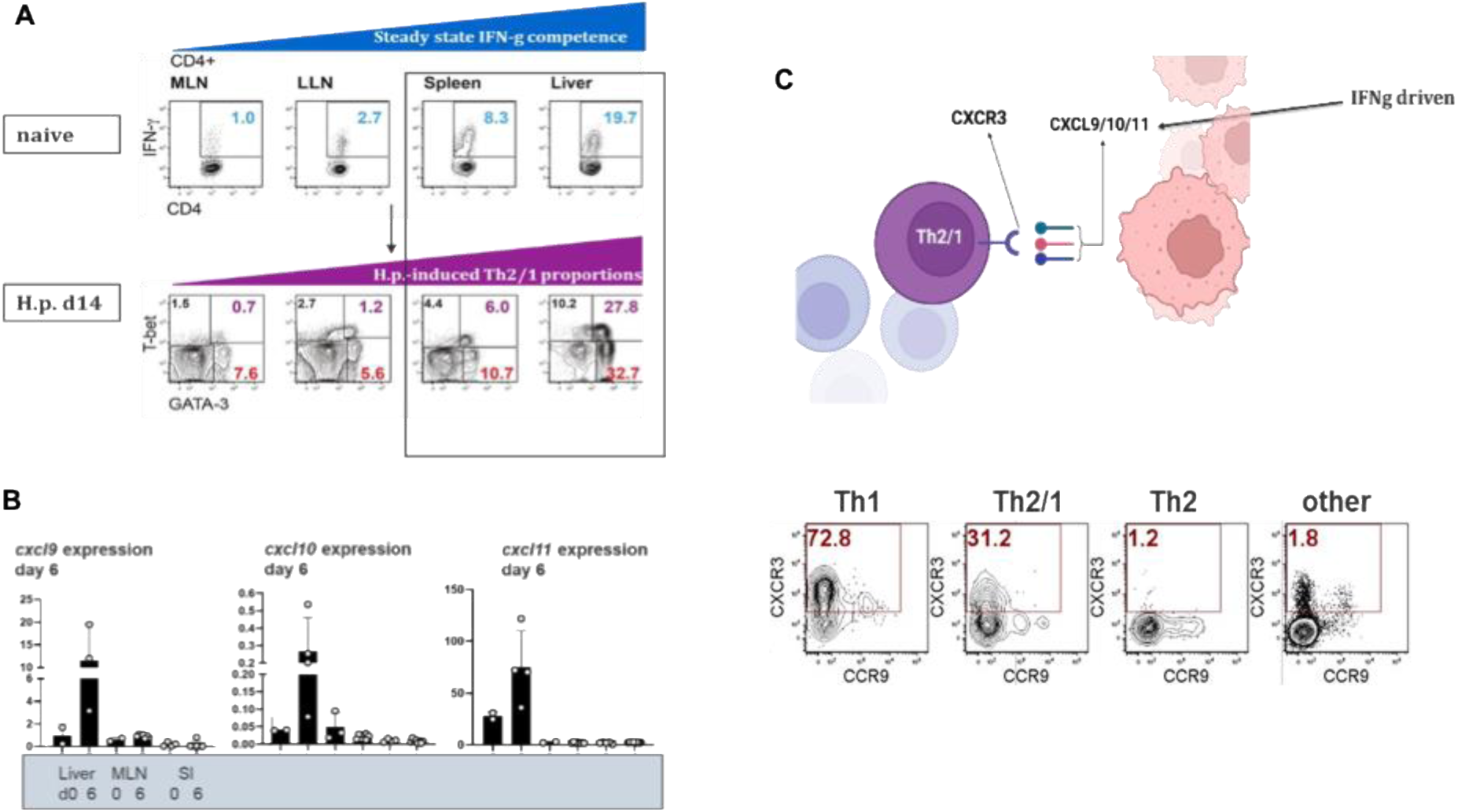
Steady-state IFN-γ drives CXCR3-mediated tissue accumulation of Th2/1 cells. (A) *Top:* Baseline IFN-γ expression in CD4+ T cells from naïve mice shows organ-specific IFN-γ competence. *Bottom:* T-bet+ vs. GATA-3+ T cell proportions at day 14 post-infection reveal infection-induced Th2/1 polarization. (B) CXCR3 expression is restricted to IFN-γ-competent Th1 and Th2/1 subsets (Liver). Plots show receptor distribution across T helper subsets, with percentages indicated. (C) qPCR analysis of CXCR3 ligand genes (*cxcl9, cxcl10, cxcl11*) in liver, MLN, and siLP at day 6 post-infection. Data show mean ± SEM.

As IFN-γ induces CXCR3 ligands (CXCL9/10/11) (J. Li et al., 2018) and Th2/1 cells express CXCR3 (Groom & Luster, 2011), we analyzed this axis. Flow cytometry revealed the highest CXCR3 expression on liver and spleen cells (Figure 3B; spleen data not shown). qPCR confirmed exclusive hepatic *cxcl9/10/11* expression at day 6 post-infection (Figure 3C), when Th2/1 cells emerge.

Since adult *H. bakeri* may cause gut perforation and microbial translocation (increased serum soluble CD14; Classon et al., 2022), we tested whether gut microbes drive hepatic Th2/1 accumulation. Antibiotic-mediated microbiota ablation did not alter hepatic Th2/1 populations versus untreated controls (Supp. Figure 2A, B), consistent with previous findings (Rausch et al., 2018). This shows microbial translocation is not required for Th2/1 recruitment, pointing to host-intrinsic mechanisms.

Collectively, IFN-γ-rich liver environments recruit Th2/1 cells via compartment-specific CXCR3-CXCL9/10/11 interactions (Cripps et al., 2012; Ye et al., 2010; Kapse et al., 2022), independent of microbial cues (Supplementary Figure 2).

#### GATA-3+ T cells in the liver are not generated de novo but accumulate from lymphoid generation sites during the infection

Next, we investigated whether liver T cells accumulate from lymphoid compartments or are induced locally by antigen-presenting cells. The liver accumulates bona fide Th2 cells at days 6 and 14 post-infection. Given the gut-liver axis, where the liver is exposed to gut-derived elements (Albillos et al., 2020; Zheng & Wang, 2021), parasite antigens from the gut could initiate a local immune response. We observed significantly elevated parasite-specific GATA-3+ CD4+ T cells in the liver on day 14 (Supp. Figure 3A), which were highly active cytokine producers (Supp. Figure 3B).

To investigate this, we infected mice and treated them with FTY720 (fingolimod) to block T cell egress from lymph nodes (Figure 4A). FTY720 treatment reduced CD4+ T cell numbers in the livers of both naïve and infected mice compared to untreated controls, ruling out de novo generation in the liver (Figure 4B). These data indicate that T cells accumulate in the liver from induction sites, though the purpose of this accumulation remains unclear.

**Figure 4:**
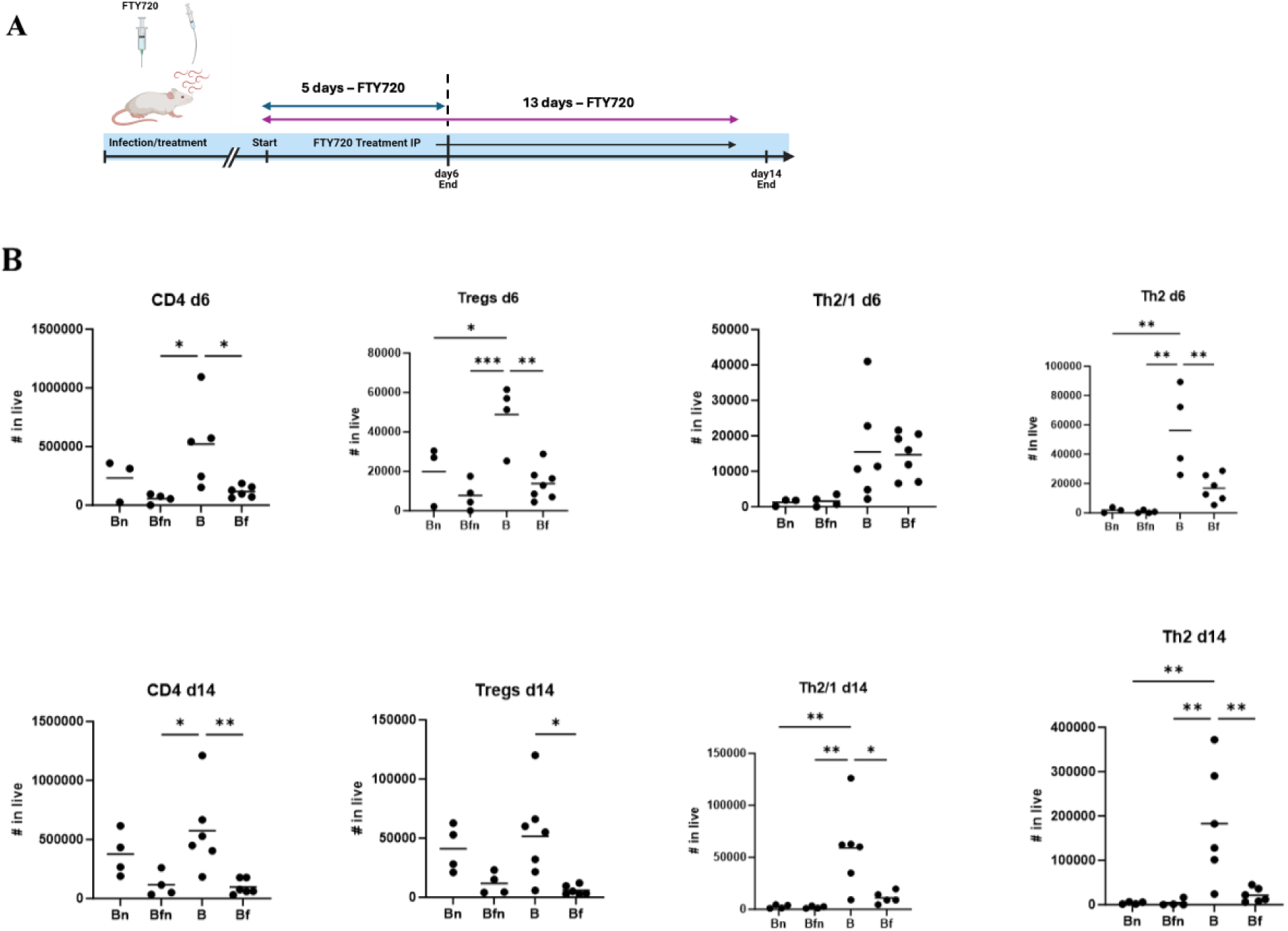
FTY720 treatment modulates hepatic T cell responses during *H. bakeri* infection. **(A)** Experimental timeline: Mice were infected with *H. bakeri* larvae and received daily intraperitoneal FTY720 injections. Liver samples were collected at days 6 and 14 post-infection. **(B)** Hepatic T cell subset counts at days 6 and 14 post-infection. Groups: Bn (naïve untreated), Bfn (naïve FTY720-treated), B (infected untreated), Bf (infected FTY720-treated). Statistical significance: *p < 0.05, **p < 0.01, ***p < 0.001 (Kruskal-Wallis with Dunn’s test or Mann-Whitney test).

#### Functional Consequences of Hepatic T Cell Accumulation During H. bakeri Infection

Our data demonstrate that the liver is a unique immunological niche during *H. bakeri* infection, accumulating T cells migrated from lymphoid organs. To investigate the functional consequences, we performed transcriptomic analysis on liver tissues from infected mice at days 6 and 14 post-infection and naïve controls (Figure 5A).

**Figure 5.**
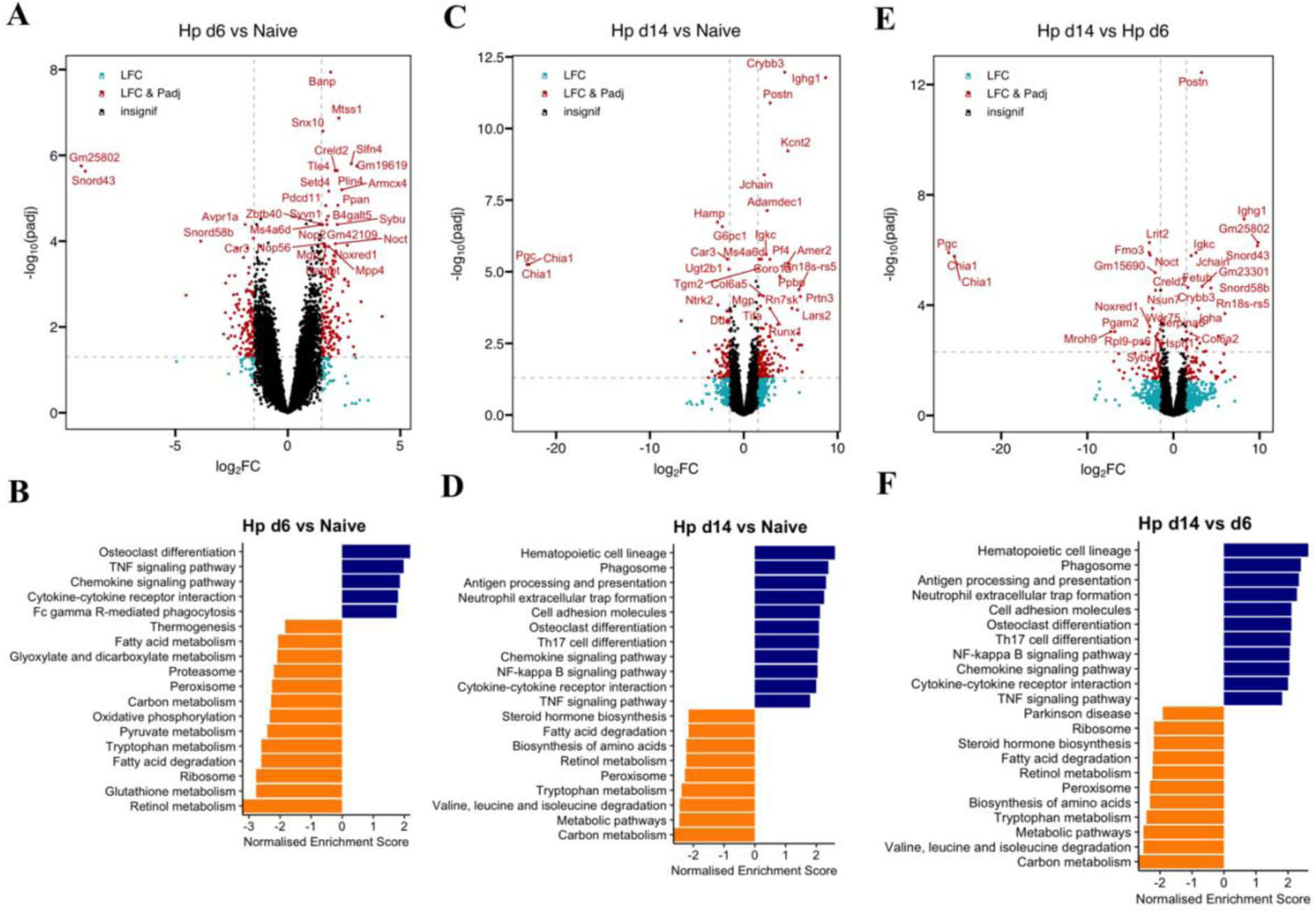
Transcriptomic and pathway enrichment analysis of hepatic response to *H. bakeri* infection. Volcano plots display differentially expressed genes (DEGs) in liver tissue for comparisons: (A) Day 6 post-infection (Hp d6) vs naïve, (C) Hp d14 vs naïve, and (E) Hp d14 vs Hp d6. Significantly regulated genes (adjusted p < 0.05, |log₂FC| > 1) are shown in red, with key genes labeled. Gene Set Enrichment Analysis (GSEA) of KEGG pathways for comparisons: (B) Hp d6 vs naïve, (D) Hp d14 vs naïve, and (F) Hp d14 vs Hp d6. Normalized Enrichment Scores (NES) show positively enriched pathways (blue) and negatively enriched pathways (orange), revealing temporal shifts in immune signaling (e.g., TNF, chemokine pathways) and metabolic processes (e.g., carbon metabolism, fatty acid degradation) during infection progression.

At day 6, differential expression analysis revealed 131 DEGs (81 upregulated, 49 downregulated; Figure 6A). This indicated concurrent immune pathway activation (TNF, JAK-STAT signaling) and suppression of metabolic pathways (oxidative phosphorylation, pyruvate metabolism), suggesting an early metabolic shift alongside immune activation (Figure 6B).

**Figure 6.**
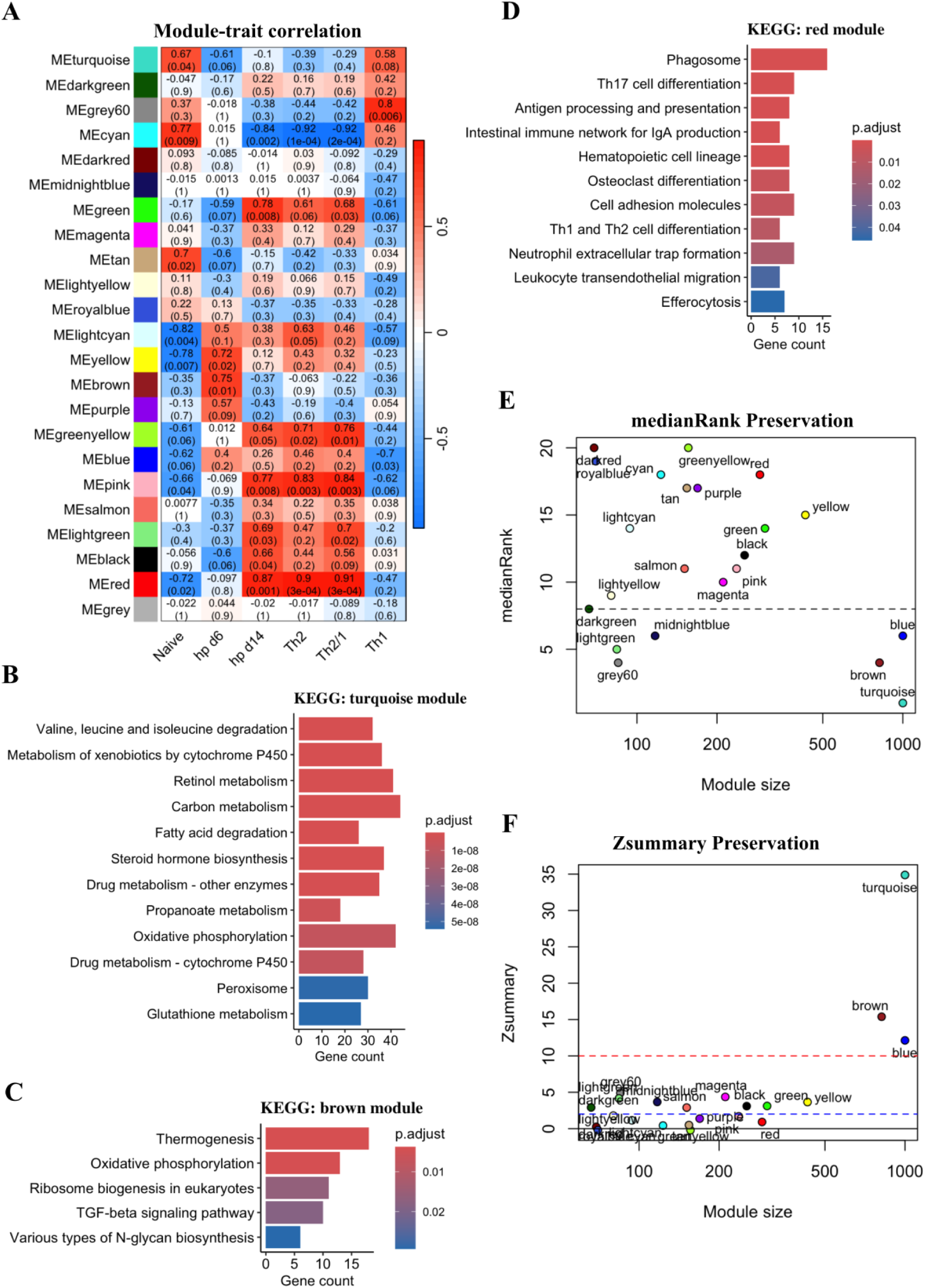
Co-expression network analysis of treatment response. (A) Module-trait correlation heatmap shows relationships between module eigengenes (colored) and experimental traits, with correlation coefficients and p-values (parentheses). (B) KEGG enrichment for the turquoise module reveals metabolic pathways (amino acid degradation, xenobiotic metabolism, oxidative phosphorylation). (C) KEGG enrichment for the red module shows immune pathways (phagosome, Th17 differentiation, antigen presentation). (D) Module preservation analysis ranks modules by medianRank (left) and Zsummary (right), distinguishing dataset-specific (grey60) from robust (turquoise) modules. (E) KEGG enrichment for the brown module identifies thermogenesis and oxidative phosphorylation pathways. Gene counts are shown; p<0.05 is considered significant. Analyses used WGCNA and KEGG databases.

By day 14, immune engagement was more pronounced, with 137 DEGs (102 upregulated, 38 downregulated; Figure 5C). Analysis showed sustained metabolic suppression but robust activation of broad (NF-κB signaling, Th17 differentiation) and anti-helminth-specific immune pathways (phagosome formation, antigen presentation) (Figure 5D). Comparing day 14 to day 6 revealed 49 DEGs (23 up-regulated, 26 down-regulated; Figure 5E). Functional analysis confirmed a shift toward active immune-related pathways and suppressed metabolism at this late stage (Figure 5F).

Overall, *H. bakeri* infection initially alters the liver’s metabolic profile, particularly reducing ATP production via oxidative phosphorylation (affecting mitochondrial processes). As infection progresses, the transcriptome shifts toward immune system activation, involving leukocyte activation and chemokine, TNF, and NF-kappa B signaling. These findings reveal stage-specific hepatic immune and metabolic reprogramming during *H. bakeri* infection.

#### Expression profiles associated with metabolism are highly preserved in the liver after blocking T cell infiltration

To determine if T cell infiltration causes the observed metabolic changes in the liver during *H. bakeri* infection, we blocked their infiltration with FTY720 and performed bulk RNA-seq on liver cells. We then conducted a comprehensive differential co-expression analysis comparing FTY720-treated and untreated datasets. Using weighted gene co-expression network analysis (WGCNA), we identified 22 distinct colour-coded gene modules in untreated liver samples (Suppl. Fig. 4A). We further assessed the association between these modules and infection status as well as immune traits (Th2, Th2/1 and Th1 responses measured by flow cytometry; see Figure 6A), revealing critical insights into the liver’s immune and metabolic responses during infection. The turquoise and cyan modules correlated with the naïve state (Figure 6A). The turquoise module was enriched for metabolic pathways like oxidative phosphorylation (Figure 6B), while the cyan module was enriched for metabolic signaling pathways (Suppl. Fig. 4B). Early infection (day 6) correlated with the brown and yellow modules (Figure 6A). The brown module was enriched for oxidative phosphorylation (Figure 6C), linking this pathway to early infection. Late infection (day 14) correlated with the red, pink, and green modules (Figure 6A). The red module also strongly correlated with Th2/1 and Th2 immune traits and was enriched for leukocyte activation and adaptive immune responses (Figure 6D), indicating a shift to immune activation by day 14.

Differential co-expression analysis between FTY720-treated and untreated groups revealed distinct module preservation. The metabolically-associated turquoise and brown modules were strongly preserved (Zsummary >10, median rank <8; Figure 6E, F). This strong preservation, alongside only six DEGs at day 6 (none linked to the brown module; Suppl. Fig. 5A), shows that early metabolic reprogramming is intrinsic and T-cell-independent (independent of the Fingolimod effect).

Conversely, the immune-associated red module was not preserved. This was confirmed by 28 DEGs at day 14, including eight red module driver genes related to B cell functions (Suppl. Fig. 5B). Collectively, these results demonstrate a dissociation between metabolic and immune reprogramming. Metabolic alterations occur independently of lymphocyte recruitment, while immune gene expression is significantly influenced by T cell infiltration.

#### Helminth-Derived Factors and Alarmins Disrupt Liver Metabolism During Early H. bakeri Infection

Our sequencing data show the liver’s metabolic profile is altered early in *H. bakeri* infection, a shift not caused by T cell infiltration (Figures 4, 5 & 6). We hypothesize this change is instead driven by *H. bakeri* excretory-secretory products (HES) or alarmins.

HES from larvae and damage-associated molecules from tissue migration may reach the liver via circulation, disrupting metabolism (Figures 5 & 6). To test this, we cultured mouse liver parenchymal and non-parenchymal cells with HES or alarmins in media containing galactose (to force mitochondrial ATP production) or glucose. Both HES and alarmins impaired cytoplasmic and mitochondrial ATP production in liver cells (Figure 7B).

**Figure 7.**
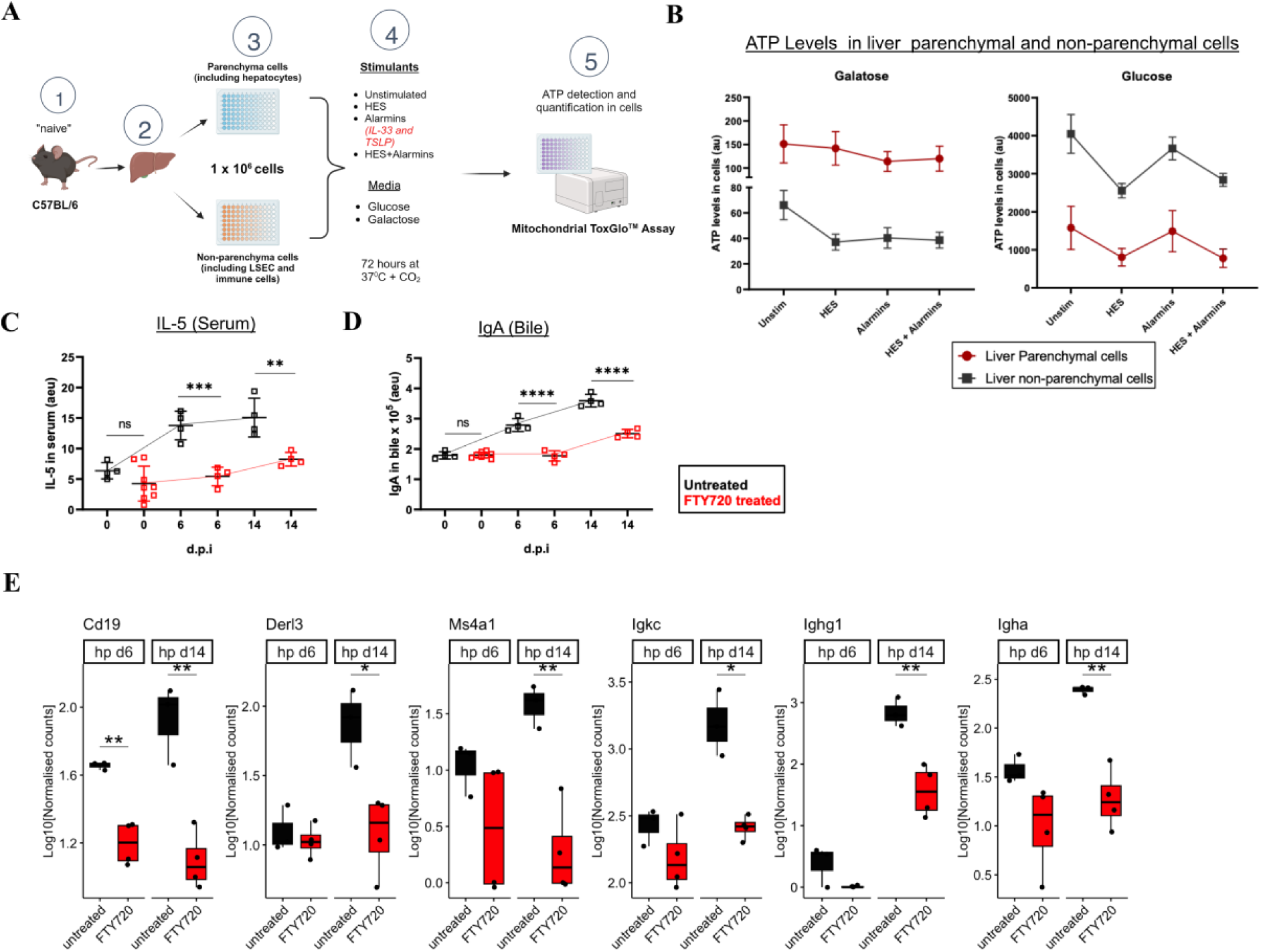
Mitochondrial function and systemic responses following alarmin and HES stimulation. (A) Experimental workflow: Liver cells from naïve C57BL/6 mice were separated into parenchymal and non-parenchymal fractions, plated at 1×10⁶ cells/well, and treated with HES, alarmins (IL-33/TSLP), or a combination of both, in glucose/galactose media for 72 hours. ATP levels were quantified using Mitochondrial ToxGlo™ Assay. (B) Total cellular ATP production from glucose (top) and galactose (bottom) metabolism in liver cells under the indicated stimulation conditions. (C) Serum IL-5 levels during *H. bakeri* infection with FTY720 treatment, showing a significant reduction post-treatment. (D) Biliary IgA levels during infection and FTY720 treatment. (E) *Liver gene* expression of *Cd19*, *Derl3*, *Ms4a1*, *Igkc*, *Ighg1*, and *Igha,* quantified in untreated and FTY720-treated mice at days 6 and 14 p.i. (*\** < 0.05, ** < 0.01). Data represent mean ± SEM.

We also found that liver T cells are highly active IL-5 producers compared to other sites (Figure 1). Because liver T cells originate from induction sites where they are not hyper IL-5 Producers, this finding indicates that they acquire this ability upon infiltration into the liver, likely as a direct result of early metabolic changes in the infected hepatic microenvironment.

Also, since IL-5 enhances IgA production and eosinophil-mediated mucosal immunity (Carreto-Benaghi et al., 2024; Yanagibashi et al., 2025), we investigated a liver-mucosal link. The accumulation of IL-5-competent T cells in the liver corresponded with increased serum IL-5 and elevated bile IgA (Figure 7C, D). Blocking T cell recruitment with FTY720 drastically reduced serum IL-5 and bile IgA (Figure 7C, D). Furthermore, FTY720 treatment significantly reduced expression of the *Igha* gene and other B cell-associated genes (*Cd19, Ms4a1, Igkc, Ighg1*) at days 6 and 14 post-infection (Figure 7E, suppl. Fig. 5C).

The coordinated reduction of *Igha, Cd19, Ms4a1, Igkc, and Ighg1* suggests attenuation of a broader B cell and class-switched humoral response. *Igha* is the most direct marker of IgA-expressing B cells, whereas *Cd19 and Ms4a1* (CD20) indicate mature B cell abundance and activation, and *Igkc* marks general immunoglobulin-producing B cells (Rickert et al., 1995; Depoil et al., 2008). *Ighg1* marks type 2-associated IgG1 class switching and can contribute to mucosal IgA responses through sequential class-switch recombination (Snapper et al., 1988; Siniscalco et al., 2025). Thus, the simultaneous reduction of these genes following FTY720 treatment is consistent with impaired B cell activation, class-switched antibody responses, and IgA-associated immunity. The accompanying reduction in serum IL-5 and bile IgA further supports this interpretation, as IL-5 promotes the differentiation and antibody secretion of IgA-committed B cells (Harriman et al., 1988).

These findings suggest that recruiting IL-5-competent T cells to the liver is pivotal for driving systemic IL-5 and enhancing biliary IgA responses, potentially contributing to anti-helminth immunity. The metabolic disruptions caused by *H. bakeri* open new avenues for further investigation of the liver during enteric infections.

## Discussion

Our study reveals novel aspects of gut-liver immunological crosstalk during *H. bakeri* infection, demonstrating previously unrecognized roles for LLNs and hepatic tissue in anti-helminth immunity. While gut-liver communication has been explored in viral infections (Brown et al., 2023; Zheng et al., 2013), this represents the first comprehensive examination during intestinal helminth infection, which affects a quarter of the world’s population. LLN enlargement despite intestinal parasite confinement suggests an active role in coordinating anti-helminth immunity rather than responding to direct parasite exposure (Adjah et al., 2025).

The finding that both MLNs and LLNs generate Th2 and T follicular helper cells expands our understanding of helminth-induced immune responses (Adjah et al., 2025). While MLNs are established sites for Th2 induction (Strandmark et al., 2017; I. L. King et al., 2010), LLN involvement suggests a more distributed model of immune activation, potentially reflecting evolutionary adaptation to disseminating intestinal parasites.

Contrary to reports of hepatic T cell anergy (Qian et al., 1997; Zheng & Tian, 2019), we found the liver serves as a reservoir for functional Th2/1 cells throughout infection, challenging traditional views of hepatic tolerogenicity (Callery et al., 1989; Jenne & Kubes, 2013). CXCR3-mediated Th2/1 recruitment, likely driven by IFN-γ and related chemokines, suggests sophisticated homing mechanisms beyond gut microbial translocation and extends earlier observations of hepatic CD4+ T cell accumulation during helminth infection (Mohrs et al., 2005; Min et al., 2004).

The functional dichotomy between hepatic and intestinal T cells reveals important mechanisms of immune adaptation. While hepatic T cells maintained proliferative capacity, intestinal counterparts showed reversible exhaustion phenotypes, supporting a model where the liver serves as an immunological reserve. The parallel peritoneal cavity response (Mohrs et al., 2005; Fodor et al., 2022) further emphasizes the importance of non-lymphoid sites in sustaining anti-helminth immunity.

Metabolic reprogramming toward glycolysis, mediated by parasite excretory-secretory products, creates a glucose-rich environment that may facilitate T cell recruitment and function. The strong association between late infection and Th2/Th2/1-related gene networks suggests that accumulating T cells modulate local and systemic immunity via the gut-liver axis. Naïve liver metabolic signatures indicate preconditioning of subsequent anti-helminth responses.

Temporal patterns show progressive reshaping of the hepatic microenvironment. Early metabolic downregulation coincides with initial immune activation, while later stages show sustained metabolic suppression with full immune engagement. This metabolic-immune interplay likely favors Th2 and Th2/1 responses, potentially explaining Th1-to-Th2/1 conversion. Enhanced cytokine production capacity following non-specific stimulation suggests the liver microenvironment potentiates T cell responsiveness. Since the liver doesn’t generate T cells de novo (Figure 4), most T cells likely acquire this capability after trafficking to the liver, a phenomenon we term ‘functional editing’.

This metabolic adaptation particularly supports hepatic IL-5 production, evidenced by reduced serum IL-5 following FTY720-mediated T cell blockade (Figure 7C). The potential liver-intestine axis linking hepatic IL-5 production and bile IgA levels has important implications for mucosal immunity. Parasite manipulation of hepatic metabolism may create a favorable microenvironment, with immune cell infiltration representing a secondary response.

The biliary immune response demonstrates functional consequences: hepatic Th2 cells rapidly acquire IL-5-producing capacity, enabling timely B cell instruction for IgA-secreting plasma cell differentiation (Figure 7D). This elevates biliary IgA levels, providing crucial protection in the upper small intestine. FTY720 experiments confirm T cell recruitment dependence, underscoring the liver’s role as a mucosal immunity coordinator despite anatomical distance from infection.

Although T cell infiltration shapes immune-related gene expression (non-preserved black module, Figure 6F), metabolic perturbations arise independently of lymphocyte recruitment, revealing dissociation between metabolic and immune reprogramming. These parasite-driven metabolic shifts may represent a host-defense strategy sustaining T cell rejuvenation and anti-helminth effector functions.

While we established parasite-specific liver T cells, their role in infection resistance/susceptibility remains unresolved. Future adoptive transfer experiments could elucidate functional contributions to infection outcomes and memory phenotype assessment for long-term protective immunity.

Several questions remain: mechanisms of *H. bakeri* antigen hepatic delivery, the functional significance of the liver-PEC-siLP axis in parasite clearance/persistence, and the clinical implications of metabolic modulation in veterinary/human contexts.

In conclusion, our study redefines the liver’s role in helminth infection as a dynamic organ that serves as both a reservoir and an activation site for effector Th2/1 cells while integrating metabolic and immune responses. The liver’s high IFN-γ environment, CXCR3-mediated T cell recruitment, and unique metabolic adaptations collectively support its critical immunological function, sustaining immune responses across chronic and memory infection phases (Figure 8). These findings open new avenues for understanding chronic helminth infections and developing interventions exploiting hepatic immunological and metabolic properties. Future research should focus on the molecular mechanisms of metabolic-immune crosstalk, therapeutic modulation strategies, and harnessing these pathways to improve vaccine efficacy or parasite clearance.

**Figure 8:**
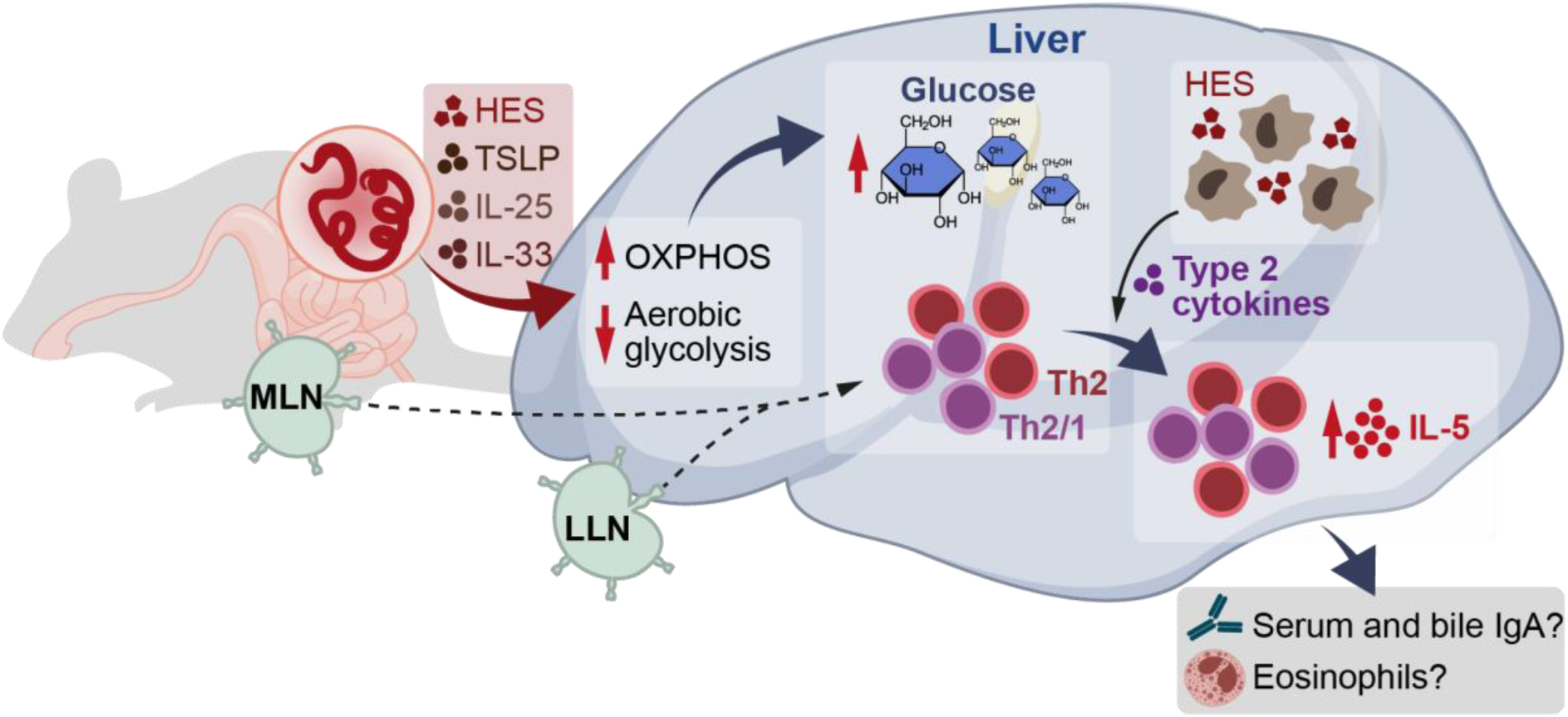
Graphical abstract. During early *H. bakeri* infection, parasite antigens and alarmins reach the liver, downregulating oxidative phosphorylation (OXPHOS) genes and promoting aerobic glycolysis. This metabolic shift triggers hepatic glycogen breakdown, creating a glucose-rich microenvironment that attracts T cells and enhances their effector function (robust IL-5 production) through “functional editing.” Concurrently, these T cells respond to translocated parasite antigens, potentially protecting the liver from hyperinflammation.

## Methods

### Mice, Infections, and Treatment

All experiments complied with German animal protection guidelines and were approved by the relevant ethics committee (LAGeSo G0176/20, 2026-093-G). Female BALB/c and C57BL/6 mice (Janvier Labs) were infected with 200 *H. bakeri* L3 larvae by oral gavage (Behnke et al., 2009). For specific experiments, BALB/c mice received intraperitoneal recombinant IFN-γ (2.5 µg) twice daily for five days (Affinass et al., 2018), an antibiotic cocktail in drinking water starting 7 days pre-infection (Rausch et al., 2018), or FTY720 (5 mg/kg) (Classon et al., 2022). Mice were anesthetized with xylazine/ketamine as required and euthanized by cervical dislocation.

### Cell Isolation

Blood was collected into FACS buffer. Mesenteric lymph nodes (MLN), liver-draining lymph nodes (LLN), peritoneal exudate cells (PEC), livers, and spleens were processed into single-cell suspensions using 70 µm strainers (BD Biosciences). Tissues were washed in RPMI (1% FCS, penicillin/streptomycin); splenocytes underwent RBC lysis. Small intestinal lamina propria (siLP) and liver lymphocytes were isolated via enzymatic digestion and Percoll gradient separation (Yordanova et al., 2022).

### Flow Cytometry

Cells were stained for homing markers CCR9 (clone CW-1.2; PE-Cy7), α4β7 (clone DATK32; biotin), and CXCR3 (clone CXCR3-173; BV421) at 37°C for 30 min (Adjah et al., 2024), followed by viability dye (eFluor506/780) incubation. Blood samples underwent erythrocyte lysis (BD FACS). Surface CD4 staining (clone RM4-5) was performed before fixation. Intracellular staining for FoxP3 (clone FJK-16s), GATA-3 (clone TWAJ), T-bet (clone 4B10), and Ki-67 (clone SolA15) used specific antibodies (Kapse et al., 2022). Dendritic cells were identified with MHC II (clone M5/114.15.2), CD11c (clone N418), and CD103 (clone 2E7) antibodies. For cytokine detection, cells were stimulated with PMA/ionomycin and Brefeldin A before staining (Steinfelder et al., 2017). Apoptosis was assessed using Annexin V kits (Thermo Fisher). Fc receptors were blocked with anti-CD16/32 (clone 93). All antibodies were from BioLegend, Thermo Fisher, or BD Biosciences. Data were acquired on FACSCanto II or FACSAria III instruments (BD Biosciences) and analyzed with FlowJo (Tree Star).

### Cell Culture and Stimulation

Cells from MLN, LLN, spleen, PEC, liver, and siLP were cultured in 96-well plates with RPMI/10% FCS and stimulated with PMA/ionomycin (1 µg/ml) for 3-4 hours at 37°C/5% CO₂ before flow cytometric analysis (Mohrs et al., 2005).

### ELISA

Parasite-specific IgA and IL-5 in serum and bile were measured by sandwich ELISA (Urban et al., 1991), with absorbance read at 450 nm (BioTek Synergy H1).

### Mitochondrial Function Assay

Liver parenchymal and non-parenchymal cells were plated at 200,000 cells/ml in glucose- or galactose-supplemented media. After 90-minute compound incubation, mitochondrial function was assessed using the Mitochondrial ToxGlo™ Assay (Promega), measuring fluorescence (485/520-530nm) and luminescence.

### Gene Expression Analysis

RNA from intestinal samples was extracted (InnuPrep RNA Mini Kit; Analytik Jena), reverse transcribed (High-Capacity RNA-to-cDNA kit; Applied Biosystems), and amplified using SYBR Green I Master Mix with primers for ccl25, cxcl9, cxcl10, and cxcl11. Expression was normalized to GAPDH and calculated via the 2-ΔΔCT method (Livak & Schmittgen, 2001).

### RNA Sequencing and Analysis

Total liver RNA was extracted (innuPREP DNA/RNA Mini Kit; AJ Innuscreen), and libraries were prepared from polyA-enriched RNA (NEBNext kits; NEB). Sequencing was performed on Illumina NovaSeq 6000 (PE 150). Reads were quality-checked (FastQC; Andrews, 2010), trimmed (fastp; Chen et al., 2018), and aligned to mm39 (STAR; Dobin et al., 2013). Gene counts were obtained (featureCounts; Liao et al., 2014), and differential expression analysis used DESeq2 (|log₂FC| ≥ 1, padj < 0.05; Love et al., 2014). Weighted gene co-expression network analysis (WGCNA) identified modules from untreated samples (Langfelder & Horvath, 2008). Functional enrichment was performed with clusterProfiler (Yu et al., 2012).

### Statistics

Data were analyzed in GraphPad Prism. Normality was assessed (Shapiro-Wilk); group comparisons used ANOVA/Kruskal-Wallis with Tukey’s/Dunn’s tests, or t-test/Mann-Whitney test for two groups.

## Author Contributions

JA, SR, SH: study design. JA: experiments. JA, ZDM, SR: data analysis and interpretation. JA, ZDM, SR, SH: manuscript writing. ZDM, AM, SR, SH: manuscript review.

## Acknowledgments

We thank Y. Weber, B. Sonnenburg, B. Anders, M. Müller, C. Palissa, and F. Möbus for technical assistance. We also thank Dr. A. Winkler for creating the graphical abstract.

## Data Availability

All data are included in the manuscript/supplementary materials or available from the corresponding author upon request.

