## Supplementary Figure 1 for "Liver-Resident Metabolic Reprogramming and Th2/1 Cell Accumulation Drive Anti-Helminth Immunity During *H. bakeri* Infection"

Running title: **Liver metabolic-immune crosstalk in Helminth infection**

Authors

Joshua Adjah^1*^, Zaneta D. Musimbi^1,2#^,3^#^, Ankur Midha^1^, Susanne Hartmann^1^, Sebastian Rausch^1^*

**Affiliations**

1 Institute of Immunology, Department of Veterinary Medicine, Freie Universität Berlin, 14163 Berlin, Germany

2 Kenya Medical Research Institute (KEMRI) - Wellcome Trust Research Programme, 80108 Kilifi, Kenya

3 Pwani University Biosciences Research Centre (PUBReC), Pwani University, 80108 Kilifi, Kenya.

\# Current affiliation

**Corresponding Author:**

Joshua Adjah

Institute of Immunology

Department of Veterinary Medicine, Freie Universität Berlin

14163 Berlin, Germany

**SUPPLEMENTARY DATA**

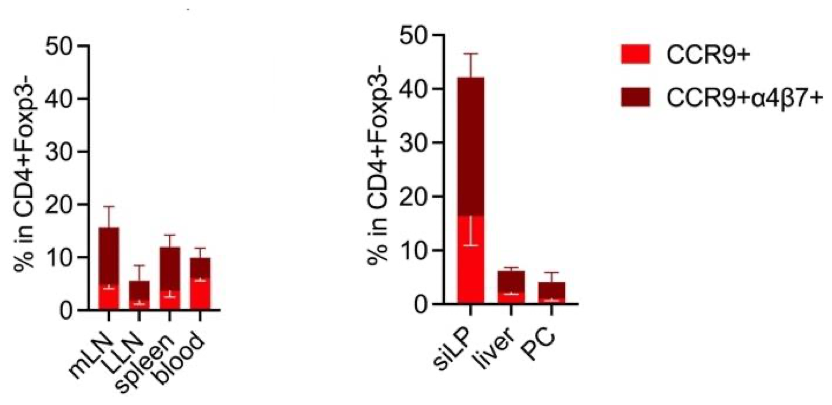

**Supp. Figure 1:** **Frequency of CCR9+ and CCR9+α4β7+ T cell subsets in CD4+Foxp3− T cells.** Bar graphs depict the percentage of CCR9+ and CCR9+α4β7+ cells within the CD4+Foxp3− T cell population, in MLN, LLN, Spleen, Blood, siLP, Liver, and PC. Error bars represent the standard error of the mean. Data are representative of at least two independent experiments.
