## Supplementary Figure 2 for "Liver-Resident Metabolic Reprogramming and Th2/1 Cell Accumulation Drive Anti-Helminth Immunity During *H. bakeri* Infection"

Running title: **Liver metabolic-immune crosstalk in Helminth infection**

Authors

Joshua Adjah^1*^, Zaneta D. Musimbi^1,2#^,3^#^, Ankur Midha^1^, Susanne Hartmann^1^, Sebastian Rausch^1^*

**Affiliations**

1 Institute of Immunology, Department of Veterinary Medicine, Freie Universität Berlin, 14163 Berlin, Germany

2 Kenya Medical Research Institute (KEMRI) - Wellcome Trust Research Programme, 80108 Kilifi, Kenya

3 Pwani University Biosciences Research Centre (PUBReC), Pwani University, 80108 Kilifi, Kenya.

\# Current affiliation

**Corresponding Author:**

Joshua Adjah

Institute of Immunology

Department of Veterinary Medicine, Freie Universität Berlin

14163 Berlin, Germany

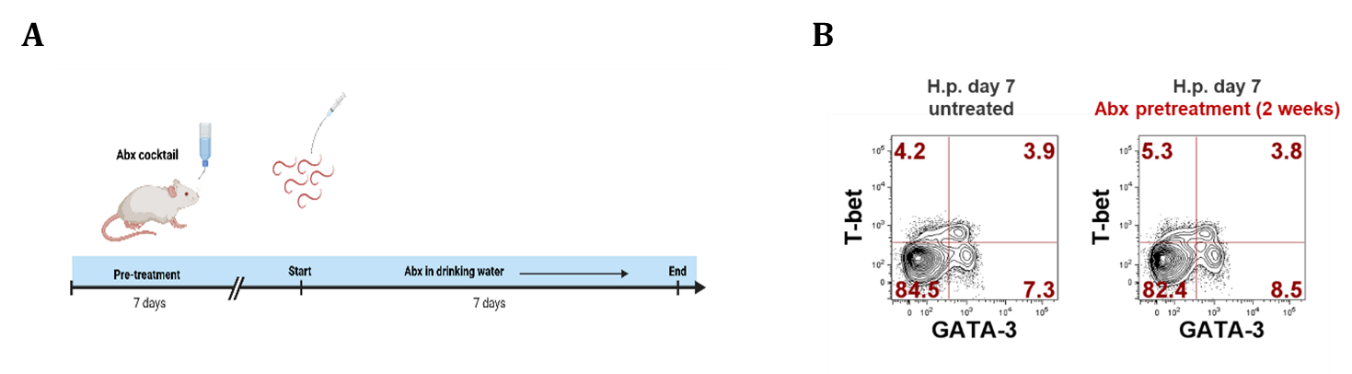

**Supp. Figure 2. Antibiotic pretreatment does not alter hepatic Th2/1 cell proportions during H. bakeri infection.** **(A)** Experimental timeline: Mice received an antibiotic cocktail in drinking water for 7 days pre-infection, continuing through 7 days post-infection before sacrifice. **(B)** Flow cytometry analysis of T-bet and GATA-3 expression in hepatic CD4+ T cells at day 7 post-infection, comparing untreated and antibiotic-pretreated groups. Quadrant percentages show Th1, Th2, and Th2/1 subset frequencies.
