## Supplementary Figure 3 for "Liver-Resident Metabolic Reprogramming and Th2/1 Cell Accumulation Drive Anti-Helminth Immunity During *H. bakeri* Infection"

Running title: **Liver metabolic-immune crosstalk in Helminth infection**

Authors

Joshua Adjah^1*^, Zaneta D. Musimbi^1,2#^,3^#^, Ankur Midha^1^, Susanne Hartmann^1^, Sebastian Rausch^1^*

**Affiliations**

1 Institute of Immunology, Department of Veterinary Medicine, Freie Universität Berlin, 14163 Berlin, Germany

2 Kenya Medical Research Institute (KEMRI) - Wellcome Trust Research Programme, 80108 Kilifi, Kenya

3 Pwani University Biosciences Research Center (PUBReC), Pwani University, 80108 Kilifi, Kenya.

\# Current affiliation

**Corresponding Author:**

Joshua Adjah

Institute of Immunology

Department of Veterinary Medicine, Freie Universität Berlin

14163 Berlin, Germany

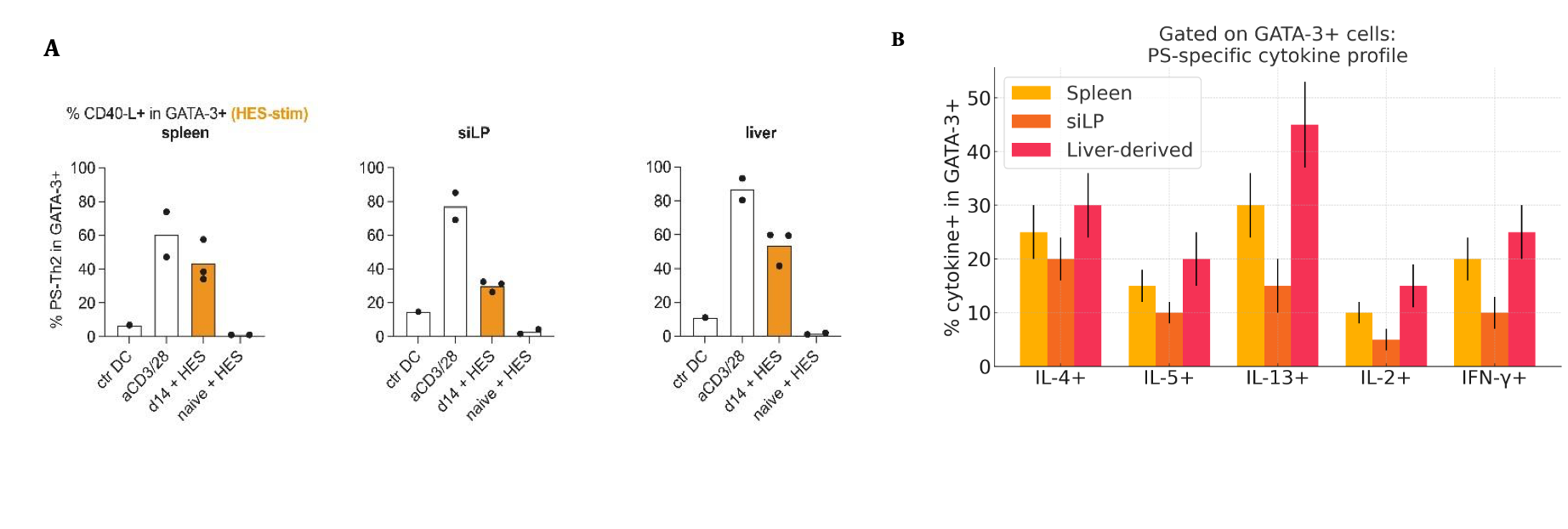

**Supp. Figure 3: Parasite-specific GATA-3+ Th2 cells and cytokine profiles in spleen, siLP, and liver. (A)** Bar graphs show the percentage of **CD40L+ cells** within the **GATA-3+** (Th2) population in the **spleen**, **liver**, and siLP. **(B)** Cytokine profiles of **GATA-3+** cells, displaying the percentage of cells producing **IL-4, IL-5, IL-13, IL-2, or IFN-γ** in response to **PS-specific stimulation of Spleen-, siLP-, and liver-derived cells**.
