## Supplementary Figure 4 for "Liver-Resident Metabolic Reprogramming and Th2/1 Cell Accumulation Drive Anti-Helminth Immunity During *H. bakeri* Infection"

Running title: **Liver metabolic-immune crosstalk in Helminth infection**

Authors

Joshua Adjah^1*^, Zaneta D. Musimbi^1,2#^,3^#^, Ankur Midha^1^, Susanne Hartmann^1^, Sebastian Rausch^1^*

**Affiliations**

1 Institute of Immunology, Department of Veterinary Medicine, Freie Universität Berlin, 14163 Berlin, Germany

2 Kenya Medical Research Institute (KEMRI) - Wellcome Trust Research Programme, 80108 Kilifi, Kenya

3 Pwani University Biosciences Research Center (PUBReC), Pwani University, 80108 Kilifi, Kenya.

\# Current affiliation

**Corresponding Author:**

Joshua Adjah

Institute of Immunology

Department of Veterinary Medicine, Freie Universität Berlin

14163 Berlin, Germany

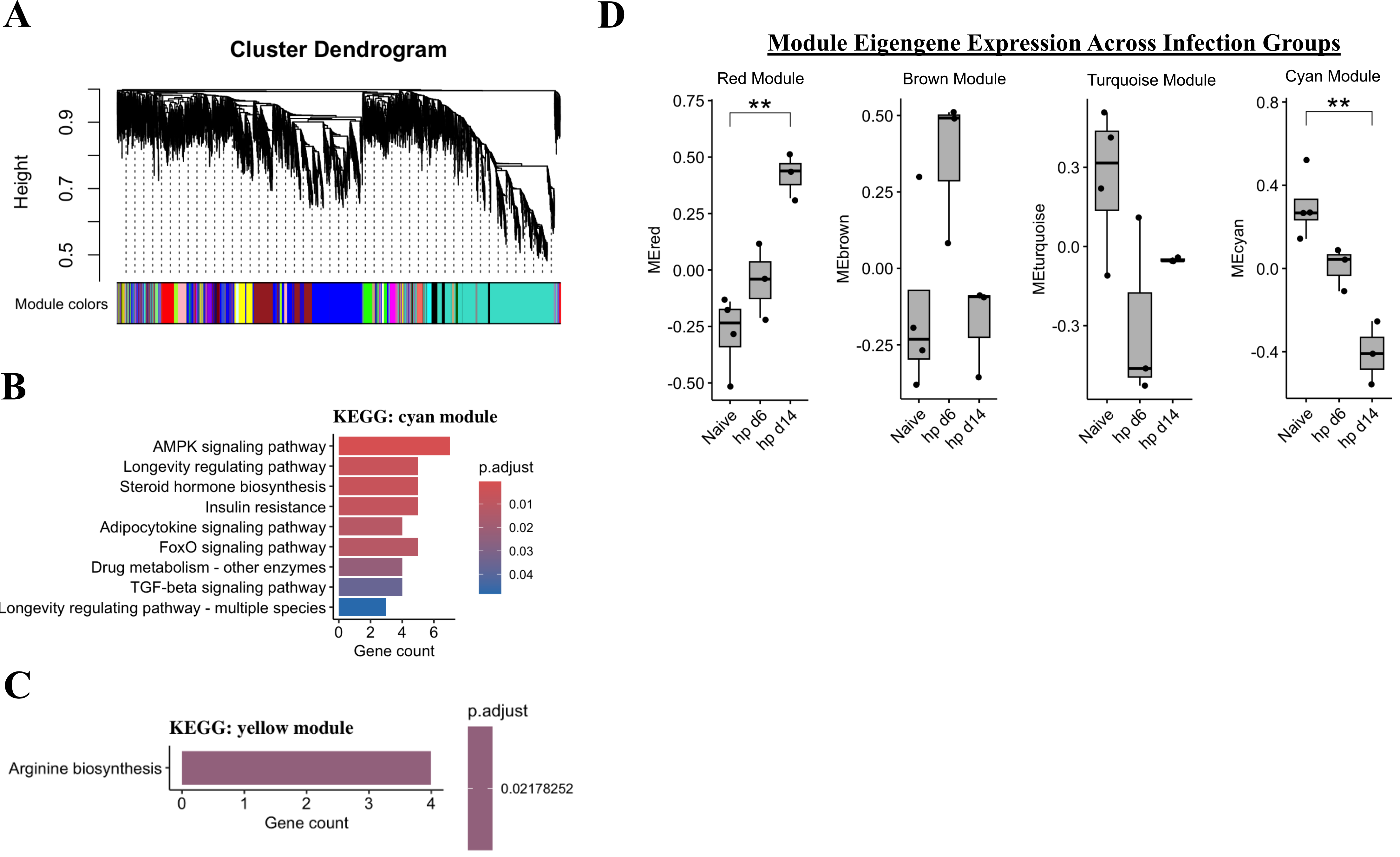

**Supp. Figure 4: WGCNA identifies infection-associated gene co-expression modules.** **(A)** Gene cluster dendrogram with module color assignments based on topological overlap. **(B)** KEGG enrichment for the cyan module shows AMPK signaling, longevity regulation, and insulin resistance pathways (adjusted p-values color-coded). **(C)** KEGG enrichment for the yellow module reveals the arginine biosynthesis pathway. **(D)** Module eigengene expression across infection groups (Naïve, hp d6, hp d14) for Red, Brown, Turquoise, and Cyan modules. Significant differences (**p<0.005, Wilcoxon test) in Red and Cyan modules indicate infection-responsive regulation.
