## Supplementary Figure 5 for "Liver-Resident Metabolic Reprogramming and Th2/1 Cell Accumulation Drive Anti-Helminth Immunity During *H. bakeri* Infection"

Running title: **Liver metabolic-immune crosstalk in Helminth infection**

Authors

Joshua Adjah^1*^, Zaneta D. Musimbi^1,2#^,3^#^, Ankur Midha^1^, Susanne Hartmann^1^, Sebastian Rausch^1^*

**Affiliations**

1 Institute of Immunology, Department of Veterinary Medicine, Freie Universität Berlin, 14163 Berlin, Germany

2 Kenya Medical Research Institute (KEMRI) - Wellcome Trust Research Programme, 80108 Kilifi, Kenya

3 Pwani University Biosciences Research Center (PUBReC), Pwani University, 80108 Kilifi, Kenya.

\# Current affiliation

**Corresponding Author:**

Joshua Adjah

Institute of Immunology

Department of Veterinary Medicine, Freie Universität Berlin

14163 Berlin, Germany

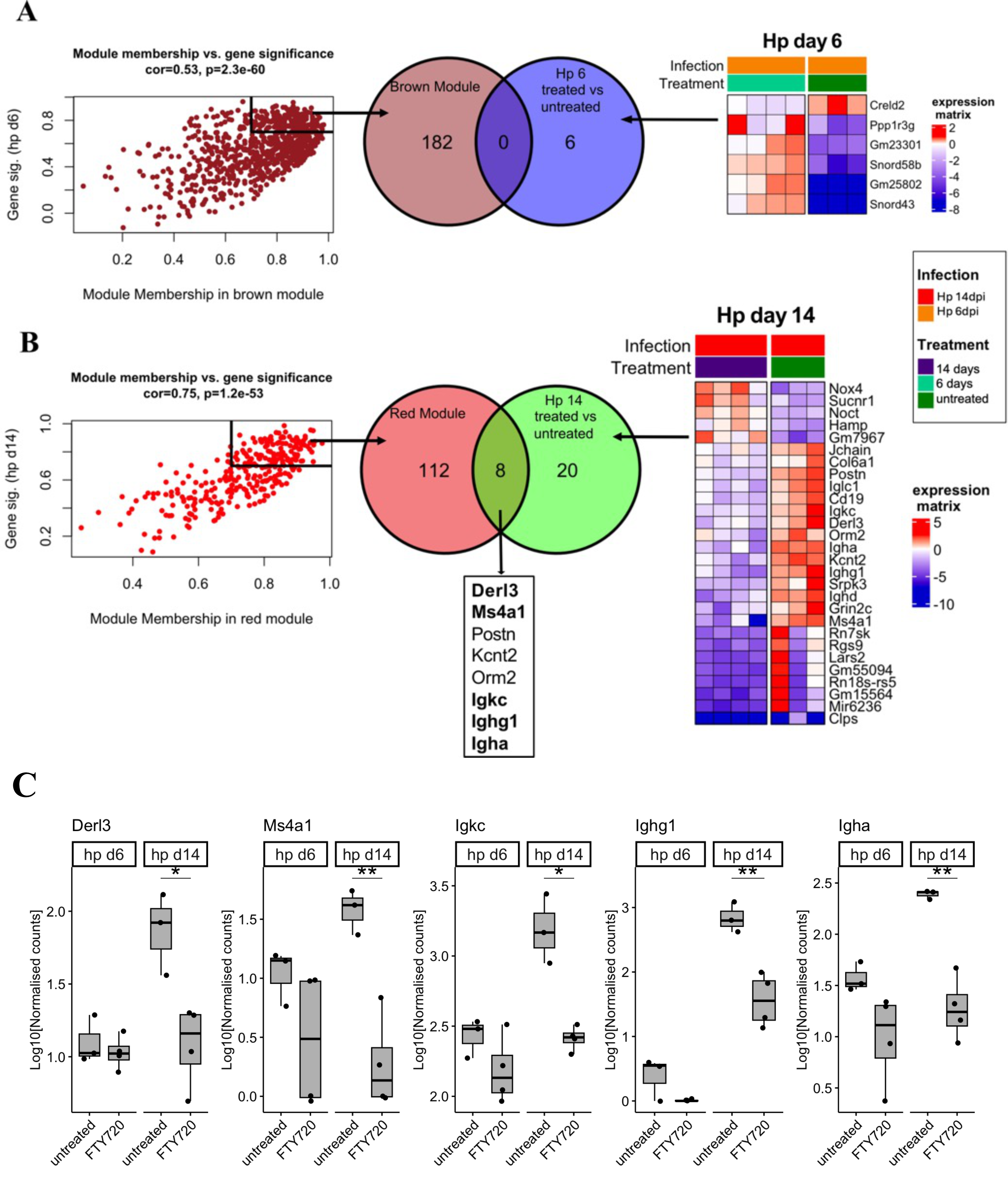

**Supp. Figure 5: Gene co-expression modules and key infection-responsive genes.** **(A)** Left: Module membership-gene significance correlation for brown module at day 6 post-infection (cor=0.53, p=3.2e-60). Middle: Overlap between brown module drivers and DEGs at hp d6. Right: Expression heatmap of overlapping driver genes across groups. **(B)** Left: Module membership-gene significance correlation for red module at day 14 (cor=0.75, p=1.2e-53). Middle: Overlap between red module drivers and DEGs at hp d14, highlighting Derl3, Ms4a1, Postn, Kcnt2, Orm2, Igkc, Ighg1, and Igha. Right: Expression heatmap of selected overlapping genes. Driver genes are defined as genes with significance >0.7 and module membership >0.7. **(C)** Normalized expression of key genes (Derl3, Ms4a1, Igkc, Ighg1, Igha) at hp d6 and hp d14 in treated/untreated groups. Significance: *p<0.05, **p<0.005, ***p<0.0005 (Wilcoxon test).
